# Complex genetic and epigenetic regulation deviates gene expression from a unifying global transcriptional program

**DOI:** 10.1101/390419

**Authors:** Mónica Chagoyen, Juan F Poyatos

**Author notes:** (MC), (JFP).

## Abstract

Environmental or genetic perturbations lead to gene expression changes. While most analyses of these changes emphasize the presence of qualitative differences on just a few genes, we now know that changes are widespread. This large-scale variation has been linked to the exclusive influence of a global transcriptional program determined by the new physiological state of the cell. However, given the sophistication of eukaryotic regulation, we expect to have a complex architecture of specific control affecting this program. Here, we examine this architecture. Using data of *Saccharomyces cerevisiae* expression in different nutrient conditions, we first propose a five-sector genome partition, which integrates earlier models of resource allocation, as a framework to examine the deviations from the global control. In this scheme, we recognize invariant genes, whose regulation is dominated by physiology, specific genes, which substantially depart from it, and two additional classes that contain the frequently assumed growth-dependent genes. Whereas the invariant class shows a considerable absence of specific regulation, the rest is enriched by regulation at the level of transcription factors (TFs) and epigenetic modulators. We nevertheless find markedly different strategies in how these classes deviate. On the one hand, there are TFs that act in a unique way between partition constituents, and on the other, the action of chromatin modifiers is significantly diverse. The balance between regulatory strategies ultimately modulates the action of the general transcription machinery and therefore limits the possibility of establishing a unifying program of expression change at a genomic scale.

## Introduction

The limited availability of the components of the cell expression machinery, for example, free RNA polymerases, cofactors, ribosomes, etc., creates a resource allocation problem that affects their activities. This differential allocation eventually represents a global program of regulation (1)(2)(3)(4), with some genes expressed at the cost of others. As the dosage of these components can be modulated by the growth rate at balanced growth, the influence of the global program has typically been studied by quantifying how the expression of genes responds to growth.

This led to the identification of laws that predict the expression of different cellular elements (3)(4)(5). Two broad models have been considered. In a first model (model 1, Fig. 1A, top), the impact of the global program is recognized by partitioning the genome in three minimal sectors (4)(5). One of them contains ribosomal genes whose expression increases with growth rate verifying its role in driving cell growth (6). Two other sectors comprise genes whose expression decreases or remains invariant with growth. The three-sector model describes therefore fundamental aspects of cellular economics (7) already advanced in the early work of bacterial physiologists (8), and acts as a basic constituent to investigate many subjects. For instance, modifications to this framework were advanced to explain the cost of unnecessary gene expression (4), the dependence of cellular composition with antibiotics (9), or the rationale behind some seemingly wasteful carbon utilization strategies (10).

**Figure 1.**
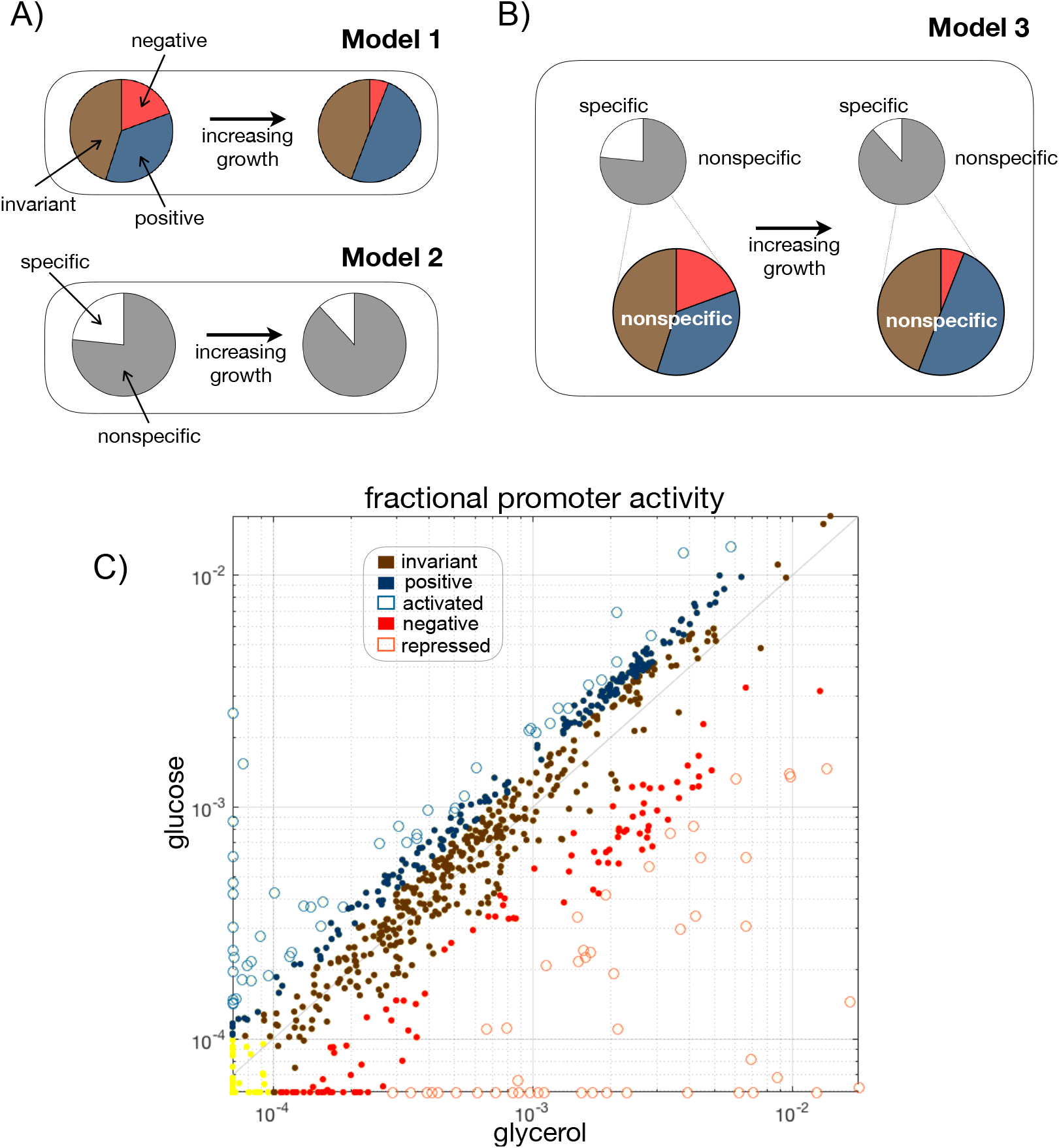
A new resource allocation model partitions the genome in five expression sectors. **A**) Top. Resource allocation model (Model 1) based on three sectors in which the expression of their constituent genes increases (positive genes, blue), decreases (negative genes, red), or remains constant (invariant genes, brown) with increasing growth rate. Bottom. Resource allocation model (Model 2) based on a specific and nonspecific sectors in which the expression of their constituent genes increases (nonspecific, white) or decreases (specific, grey) with increasing growth rate. **B**) Partition of genome expression into five sectors that combine the two previous models. A model-1-like partition appears as a fine structure within the nonspecific sector. We labelled this scheme as Model 3. **C**) Fractional promoter activity (fPA) for a transition between two example conditions increasing growth (glycerol to glucose). Promoters can be classified into the five sectors of model 3 depending on how their fPA changes (yellow dots indicate those with very low activity in both conditions). Repressed and activated promoters constitute the specific sector. Nonspecific promoters are constituted by one invariant type and two other subclasses whose fPA depends on the growing condition.

In a second model, the emphasis is not so much in the trade-offs between ribosomal genes and the rest, but between genes that follow a common pattern of expression and a minimal subset that diverges (model 2 containing two sectors: nonspecific and specific, Fig. 1A, bottom) (11). Here, the common pattern of expression is explained by a single scaling factor, which incorporates the resources of the global program that are not involved in the activation of the specific genes. This implies that most changes in expression result from a passive rather than active regulation what produces a unifying pattern that “simplifies” the expected complexities of genome-wide expression changes.

In this manuscript, we integrate both models (model 3, Fig. 1B). The new framework discriminates a fine-structure within the nonspecific sector of model 2 that corresponds to the three-sector partition representative of model 1. While the integration of both views is significant *per se*, models 1 and 2 might appear unrelated, we also show that this scheme is fundamental to help us expose how specific regulatory strategies modulate the impact of the global transcriptional program at the genome-wide level.

To this aim, we first reexamined the original data that led to model 2, i.e., promoter activity (PA) measurements of *Saccharomyces cerevisiae’s* genes obtained in different growing conditions (11), to present our new scheme. However, this data only includes a subgroup of ~900 yeast genes, and thus the quantification of the resource allocation can only be approximated. To obtain a more precise classification we considered genome-wide expression data of yeasts growing in chemostats (12) and developed a method to define the new model at a large scale. We validate our partition through functional analysis of the corresponding components.

Armed with this new framework, we focused on understanding how the global program becomes modulated by specific regulatory mechanisms, a question which is only beginning to be addressed. Indeed, within the framework of model 1 results are mixed. For instance, several recent reports in bacterial systems demonstrated the prevalence of the global expression program, while they have lowered the importance of transcription factors (TFs) controlling the assumed deviations from it. TFs seem only to fine-tune the action of the global regulation (13)(14) in combination with a few metabolites (15). More recent work in yeast argued, in contrast, about the relevance of epigenetic factors (promoter nucleosomal stability) as modulators of the global program (16).

The function of the specific regulation might appear more straightforward *a priori* in the scheme of model 2. Specific genes are indeed enriched by specific regulation, generally coupled to the particular nutrient condition, whereas the behavior of those genes within the nonspecific sector does not seem connected to a particular transcription regulation strategy (11). We thus examine within our new framework to what extent we could observe differential genetic and epigenetic regulation acting on the genes constituting each sector. We discovered well-defined regulatory patterns.

More broadly, our results put forward an integrated view of the resource allocation connected to genome-wide expression and emphasize how the global program is eventually modified by the specific regulation. Active strategies of control are certainly at work in genes within the fine-structure of the “nonspecific” response and can eventually associate the metabolic status of the cell with gene expression. Overall, this complex hierarchy of regulation in eukaryotes enhances the adjustment of genome-wide expression patterns beyond the one that could be achieved through a passive unifying program of global transcriptional control.

## Results

### Integration of two preceding resource allocation models

We first examined how PA changes with the growth rate for a subset of *Saccharomyces cerevisiae* genes. Keren *et al*. (11) presented a binary partition to describe these changes, recognizing a common response in the absolute PA values of most genes and a specific one in a much smaller subgroup. To focus on resource reallocation, we studied here fractional PA activities instead of absolute values, i.e., the fraction of PA of each gene in a given nutrient condition out of the summed activity of all genes in the set (3), and quantified their change for each pair of conditions (from low to high growth rate).

Figure 1C shows a descriptive case (glycerol to glucose conditions). The most extreme deviations of the global program correspond to the specific genes (11). Note also that the stronger allocation of “expression resources” to these genes the fewer resources to biosynthesis (reduction of the nonspecific sector) affecting growth rate (Fig. 1A, bottom). We revised the two-sector partition by further separating specific genes as specifically activated genes (fractional PA becomes much larger between conditions), and specifically repressed genes (fractional PA becomes much smaller), and also delimiting three components within the nonspecific sector: genes whose fractional PA remains approximately invariant (diagonal in Fig. 1C), positive genes, whose fractional PA moderately increases between conditions, and negative genes (fractional PA decreases to only a limited extent, see Methods for details). These three sectors relate model 2 to model 1.

We validated this new partition in two ways. First, that a set of genes follows a precise proportional response between conditions was suggested by (11) as an evidence that they share a common functionality (e.g., being part of the same pathway, etc.). Here, we find that invariant, positive and negative genes also follow a precise proportional response (Fig.S1) suggesting that our analysis identifies a biologically relevant finegrained structure. Second, we similarly expect that this fine-grained structure would parallel that proposed in model 1 (4) in terms of functional categories. This is the case. The invariant class is enriched by transcription regulation and ribosomal proteins; the latter being more extensively observed in the positive class. Indeed, positive genes are enriched by ribosomal genes (~65% of genes code for small or large subunits of the ribosome), while negative genes are enriched in ATP metabolic processes, e.g., oxidative phosphorylation or the TCA cycle (Table S1, Methods).

### The five-sector resource allocation model on a genomic scale

To substantiate the previous five-sector model based on ~900 genes, we examined a genome-wide DNA microarray dataset of yeast cells exhibiting the same range of growth rates for six limiting nutrients (12). We studied again changes of relative expression (Methods) and applied singular value decomposition (SVD) to each nutrient separately. The first and second SVD components (Fig. 2A) explain >90% of the variance in each condition (the components exhibited an analogous trend in all nutrients, Fig. S2). As a result, the fractional expression response to growth of each gene can be approximated by the linear combination of these two components (Fig. 2B).

**Figure 2.**
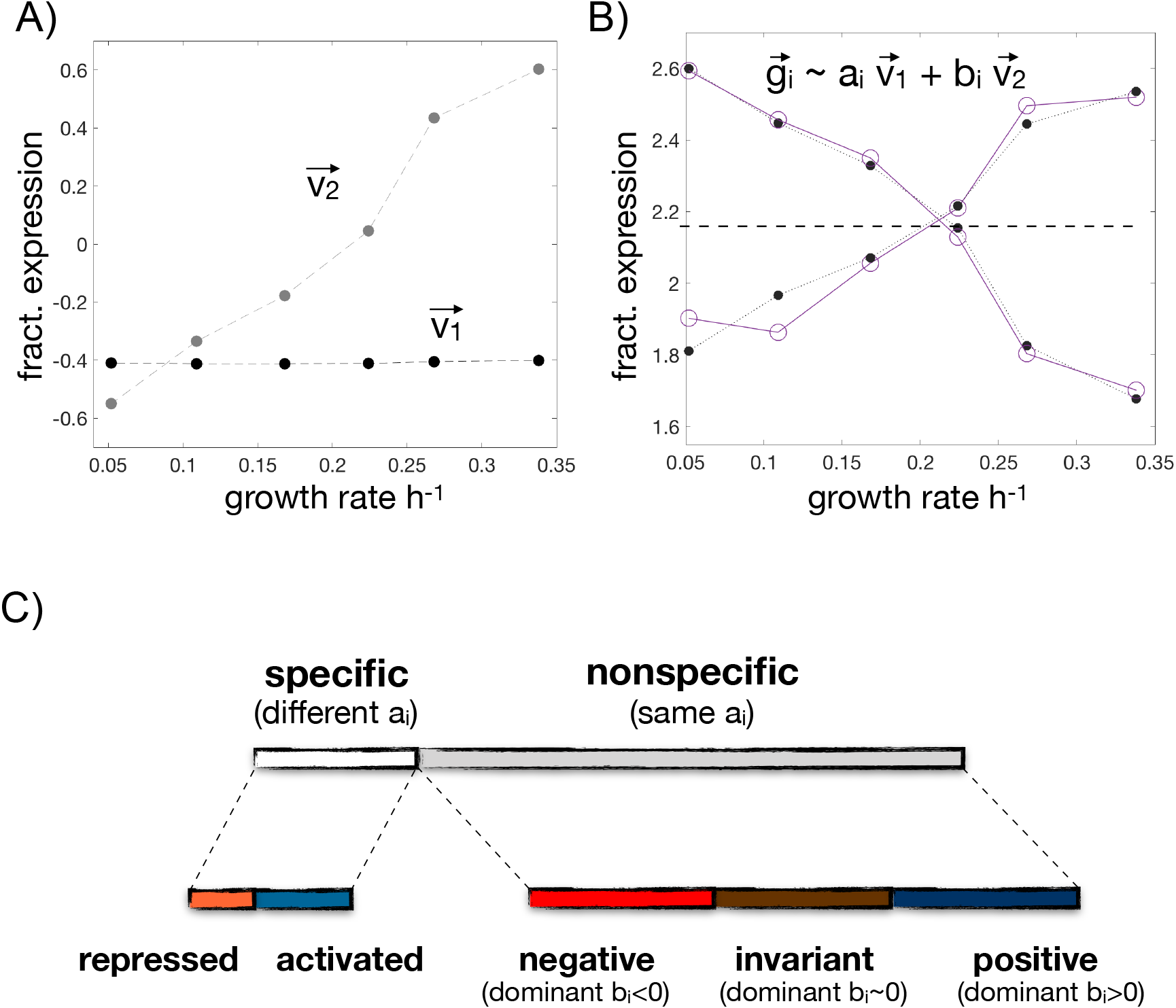
Genome-wide partition of gene expression in five sectors based on SVD. **A**) SVD components 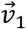 and 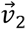 describe baseline fractional expression and dependence with growth, respectively, and together explain most of the expression variance. **B**) The fractional expression of every gene 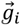 as a function of growth rate can be approximated by a linear combination of these two components, with loadings a_i_ and b_i_. We show two examples (purple circles denote the expression vector, while the black dots correspond to the two-component approximation; lines added to help visualization) with the same baseline (dashed line; same a_i_) but whose expression increases (b_i_>0) or decreases (b_i_<0) with growth. Data in **A**) and **B**) corresponds to growth in limiting glucose conditions. **C**) The baseline expression of a gene between two conditions can change (top left) or not change (top right). A gene is nonspecific if its baseline expression remains the same (similar a_i_ loadings) in >8 pairwise change of conditions (total of 15), being specific otherwise. Within nonspecific genes the second loadings b_i_’s on each nutrient condition (total of 6) enable us to determine if the gene is positive (b_i_>0), invariant (b_i_~0) or negative (b_i_<0). Specific genes can similarly be divided between repressed or activated. In this way, we can partition the genome into five sectors what generalizes the scheme discussed with PA data in a subset of yeast genes (Figure 1). See main text for details.

Furthermore, we can interpret the first element of the linear combination (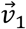) as the baseline fractional expression of the gene, which does not change with growth rate, and the second element (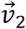) as its monotonic response to growth (Fig. 2B). This reading enables us to generalize the previous partition framework obtained with PA data. More specifically, a gene that maintains the same baseline expression (similar loading of 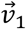 that we denoted a_i_) between two conditions involves a nonspecific response. When this response in observed in >8 pairwise changes then we reason that the gene is nonspecific (recall that there are 6 different nutrients and consequently a total of 15 pairwise nutrient changes). Genes are considered specific otherwise (Fig. 2C).

Beyond the classification above, the second component (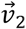) provides a quantitative score (the second loading, b_i_) to classify genes as invariant, positive or negative (b_i_~0, b_i_>0, b_i_<0, respectively; Fig. 2C. Methods). We labelled as invariant those nonspecific genes which exhibit this behavior in >3 nutrient conditions (total of 6). Nonspecific and *not* invariant genes appearing more times as positive than as negative are categorized as positive, and likewise for negative. Finally, specific genes which appear more times as positive than as negative (again in all 6 conditions) are categorized as activated, and analogously for repressed (Fig. 2C).

Finally, the functional analysis of genes within each sector agrees with that obtained with the PA data and previous reports, what substantiates the biological significance of the partition (Table S2; see also Table S1 showing how this classification maps onto the subset of genes with PA data). In this way, we have generalized at a large scale the new resource allocation model, a genome-wide classification that we use throughout the next sections.

### TFs are typically nonspecific genes

We inspect next the influence of the most direct elements related to specific regulation, i.e., TFs. But before examining their impact on the activity of the global transcription program, we asked first how TFs themselves are framed in the previous partition.

We observed that most constituent TFs of the transcriptional regulatory network (122 of a total of 133 composing the network, Methods) are nonspecific genes, i.e., they present similar basal fractional expression (a_i_ loadings) across all pairwise condition changes. Within this set, 31% presents a dominant invariant response (b_i_~0 in >3 nutrient conditions, of a total of 6), with five genes acting as invariant in all six conditions (*rsc1, mbp1, pho2, rgr1*, and *swi6*). Two of these (*mbp1, swi6*) are at the top of the network hierarchy (being involved in the mitotic cell cycle), and two are elements of relevant complexes that interact with RNA polymerase II (*rsc1* of the RSC chromatin complex, and *rgr1/med14* of the mediator complex); they can be considered as elements of a general transcriptional machinery, for which maintaining its concentration invariant across conditions could be essential. Moreover, 32% of nonspecific TFs are dominantly negative, and only 4% positive. Of note, some of the TFs whose expression decreases with growth (b_i_<0) are positive regulators of transcription in response to stress (e.g., *bur6, gcn4, rpn4*) what justifies their overexpression at low growth rates.

### Genes within each partition sector present different degree of transcriptional regulation

Is the extent of regulation of target genes dependent on which sector they belong to? We computed the mean number of regulators acting on genes within each sector. Nonspecific genes are less regulated, on average, by TFs than specific ones as expected [by 3.09 TFs *vs*. 5.06 TFs, p = 1.20 10^−4^, two-sample Kolmogorov-Smirnov (KS) test]. Within nonspecific genes, invariant genes are less regulated than nonspecific and *not* invariant ones (by 2.56 TFs *vs*. 3.3 TFs, p = 8.16 10^−13^, two-sample KS test). Finally, nonspecific and positive genes are slightly more regulated than nonspecific and negative genes (by 3.33 TFs *vs*. 3. 27 TFs, p = 0.0018, two-sample KS test). Overall, specific genes are subjected to more regulation (larger number of TFs) (Fig. 3A), while nonspecific and invariant ones show the least.

**Figure 3.**
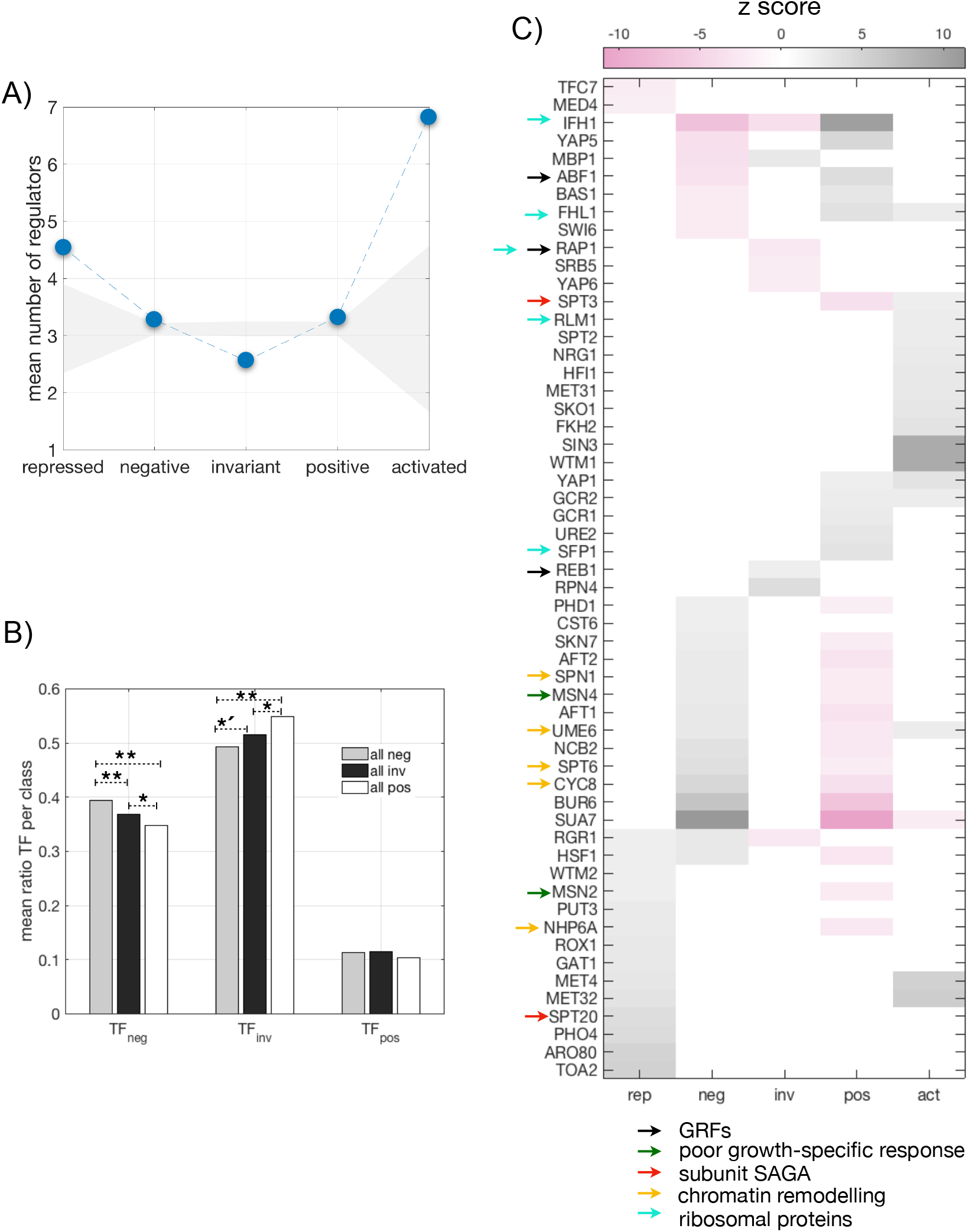
Differential integration of transcriptional regulation and the five-sector resource allocation model. **A**) Mean number of regulators acting on genes as a function of their response to growth (blue dots; dashed line to help visualization). Grey shading denotes the average null values +/- 2 standard deviations obtained by randomization. **B**) Mean ratio of the fraction of TFs of a given class with respect to growth (e.g., TF_neg_ denotes TFs which are negative genes) for each group of target genes (also for a given class; here we do not distinguish between nonspecific and specific). Histogram obtained in glucose conditions, see also Fig. S5 for other nutrients (** p < 0.001, *′ p < 0.01, * p < 0.05, twosided KS test). **C**) Regulators that act dominantly, or secondarily, in genes showing a significant regulatory coherence. Color denotes z-score with respect to a null obtained by randomization (positive values denoting enrichment; negative values denoting lack of it). Properties of some regulators are also included (arrows). See main text for details.

### TFs and their regulated genes frequently exhibit the same response to growth

Although Fig. 3A shows how the number of transcriptional interactions is reflected differentially in the sectors of the partition it does not assure us whether these interactions are functioning, e.g., regulation by TFs have been shown to be “inactive” in bacteria (13). For this, we examined several features.

We initially inspected if target genes presenting a particular growth response are enriched by TFs showing the very same response, as the similarity of the responses could indicate that these TFs do influence the expression of the target. We thus computed -for each target gene- the fraction of its regulators that behave as negative, invariant, or positive with the growth rate in a given condition (TF_neg_, TF_inv_, TF_pos_, respectively). Figure 3B shows the mean of these fractions for target genes which themselves show a negative, invariant, or positive response, respectively. Negative TFs are more likely to be found acting on target genes that are also negative (higher mean TF_neg_ on negative genes), while invariant (TF_inv_) and positive (TF_pos_) TFs regulate more often invariant and positive target genes, respectively (the latter signal is weaker and depends on the particular condition, Fig. S3). Thus, TFs that exhibit the same behavior as their cognate target gene tend to appear, on average, dominant on its regulation; this indicates that part of the regulatory structure is functional.

### Specific TFs regulate genes fitting to each partition sector

To further estimate the active regulatory character of TFs, we measured the correlation of the response to growth rate between any particular gene and all its cognate TFs (“regulatory coherence”, Methods). Specific genes showed stronger regulatory coherence than nonspecific ones (Fig. S4A), and remain coherent in more nutrient conditions (Fig. S4B), both results implying an existing contribution of TFs to deviate gene expression from the global program.

Moreover, Fig. 3C shows those TFs whose operation is particularly coherent per partition sector (Methods). One can identify here that TFs controlling specific genes are normally not acting on nonspecific ones [this is supported by earlier reports(17)]. Moreover, some of the TFs that work coherently on positive genes (gray color, Fig. 3C) are particularly absent in negative ones (pink color, Fig. 3C) and *vice versa*. In addition, those that control nonspecific genes are higher up in the network hierarchy (Methods). We also noted that some of these (significantly coherent) TFs are involved in chromatin remodeling (Cyc8, Ume6, Spt6, Msn4, Abf1, Msn2, Nhp6A, acting on nonspecific ones), or chromatin organization (Spt3, Spt2, Pho4, FKh2, Sin3, Spt20, Wtm2, Wtm1, Hif1, acting on specific genes). We examine epigenetic aspects next.

### The resource allocation model also reveals distinctive epigenetic regulation

To inspect the function of epigenetic control mechanisms, we first quantified the proportion of general transcription factors (GTFs) found within the set of TFs acting on a given gene (18). GTFs (Rap1, Abf1, Reb1, Cbf1, and Mcm1) usually have little intrinsic regulatory activity and together with the presence of chromatin remodelers (in particular, RSC-Remodels Structure of Chromatin) control an alleged general machinery of expression. We observed that GTFs constitute a larger and significant fraction in the regulation of positive genes, while the opposite is observed for negative ones (Fig. 4A).

**Figure 4.**
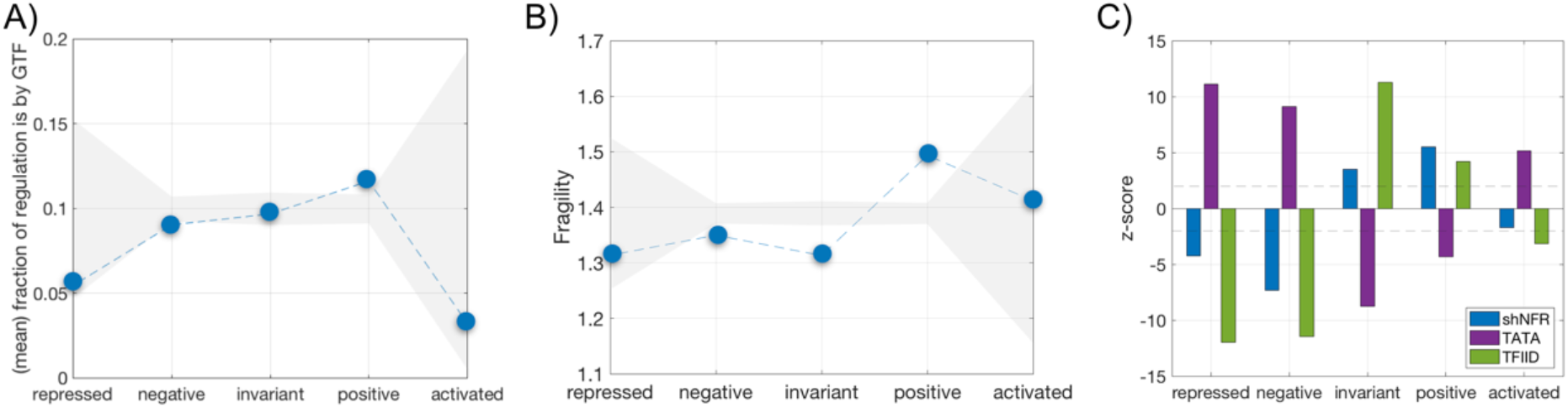
Differential integration of epigenetic regulation and the five-sector resource allocation model. **A**) Mean fraction of global transcriptional regulators (Rap1, Reb1, Cbf1 and Mcm1) within the full set of regulators acting on each gene. Grey shading denotes the average null values +/- 2 standard deviations obtained by randomization. Dashed lines to help visualization. **B**) Mean nucleosomal fragility. Shading/lines as before. **C**) Enrichment of nucleosomal free regions (shNFR, blue), presence of TATA boxes (purple), or action of TFIID global factor (green) as function of response class (measured as z-score with respect to a null by randomization; dashed line indicates z-score = +/- 2).

GTFs are also associated with particularly fragile nucleosome promoter architectures (19), a connection recently examined (16). Using this data, we computed the nucleosome landscape for the different gene classes (Methods). Promoters of positive genes are indeed enriched in fragile nucleosomes (Fig 4.B) while both negative and invariant genes lack these structures. This indicates that positive genes are more sensitive to adjust the global program of expression by means of chromatin modulation. Enrichment of other promoter features contribute to this observation (Fig 4.C), like the absence of TATA boxes, the action of TFIID over SAGA [but this precise grouping has been recently reexamined (20)], the presence of nucleosomal free regions closer to the transcriptional starting site (shNFRs) (partially associated to the previous score of fragile nucleosomes), and the dominant effect of *trans* variability (Fig. S5) (21).

### Chromatin modifiers further modulate the global program

Finally, we examined the effects of mutating different types of *trans*-acting chromatin regulators on the genes constituting the sectors using a previously assembled compendium (22) (Methods). We considered first the magnitude of change of expression (i.e., the *absolute* values of expression) before/after the mutation of several types of modifiers. With the exception of histone acetyltransferases (HATs) and TATA-binding protein related factors (TAFs), the influence of most chromatin modifiers is dominant in specific genes as compared to nonspecific ones (Fig. S6, Methods).

Within nonspecific genes, we also quantified the type of expression change (experienced after mutation of the modifiers) and found three broad relationships (Fig. 5): 1) Epigenetic regulators acting as part of a general machinery (HATs-including SAGA-, TAFs and methyltransferases) whose mutation causes a general decrease in expression, very particularly in invariant and positive classes. Indeed, work by (20) and (23) demonstrated that SAGA and TFIID are recruited to pol II promoters genome-wide and that each complex is generally required for pol II transcription, i.e., its mutation would lead to a genome-wide decrease of gene expression. 2) Regulators (e.g., histones, etc.) acting in a dual manner: increasing the expression of negative genes after mutation (remodeler as a repressor) or reducing their expression in positives (remodeler as an activator). This underlines the enrichment of negative and positive classes by stress and ribosomal genes, respectively, which are largely regulated in an opposite manner (24); a dual role of remodelers as activators and repressors have been previously reported (25)(26). And 3) regulators as broad repressors (mutation increases expression), e.g., those that represent regulation by gene silencing.

**Figure 5.**
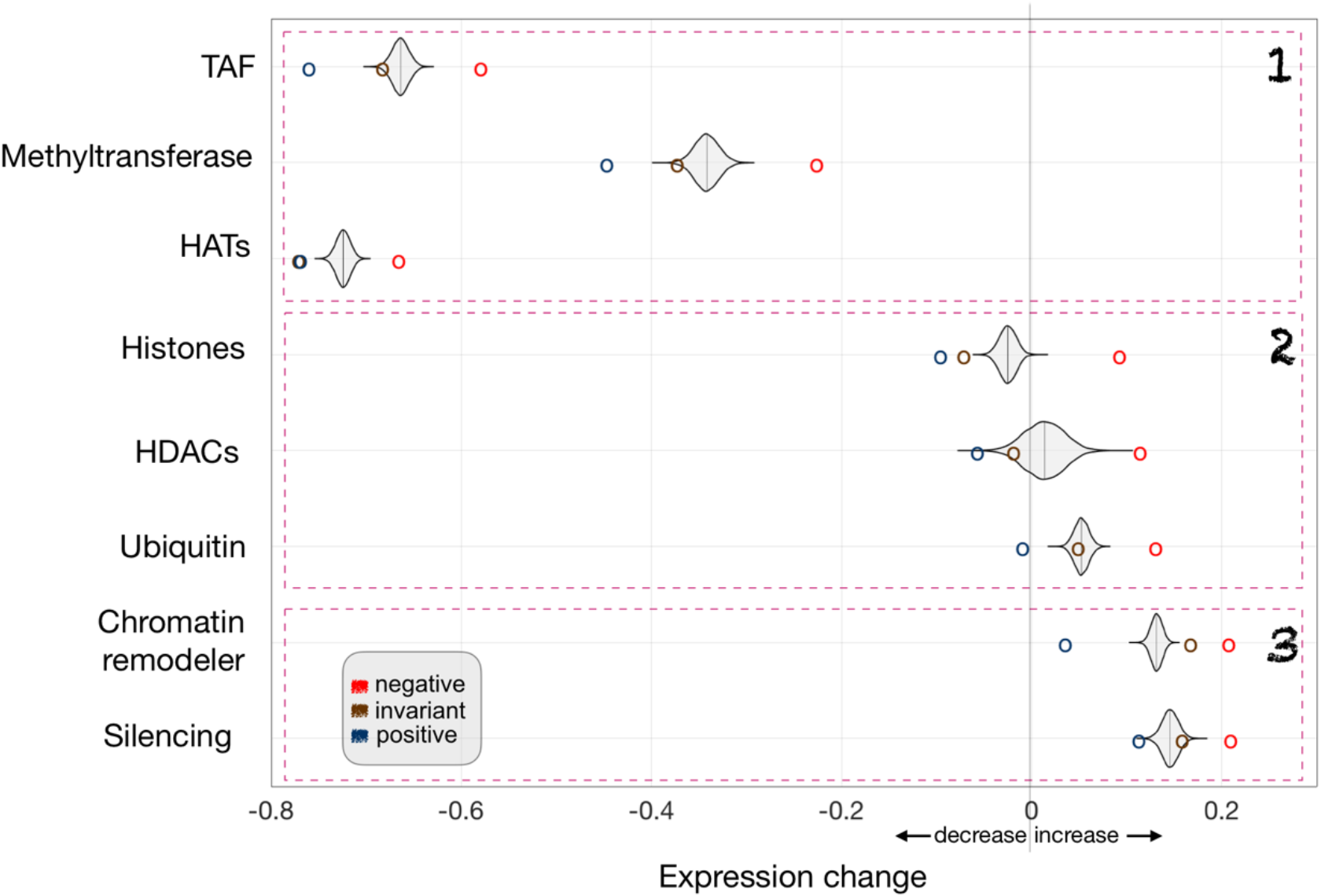
Chromatin modifiers act differentially on nonspecific genes. Mutations in chromatin modifiers reveal their diverse regulatory role. For each modifier, we plotted its mean effect, after mutation, on the genes constituting the negative (red circle), invariant (brown circle), or positive (blue circle) sectors. A kernel density plot corresponding to a null distribution obtained by permutation is also shown (Methods). We observed three broad categories: 1/ modifiers acting as activators (mutation decreases significantly the expression of invariant/positive genes), 2/ dual activator/repressor (mutation decreasing/increasing significantly expression of positive/invariant or negative genes, respectively), and 3/ repressors (mutation increasing significantly expression of negative genes). TAF: TATA-binding protein related factors; HATs: histone acetyltransferases; HDACs: histone deacetylases.

## Discussion

Recent studies emphasized how most genome-wide expression changes might be the consequence of a unifying global program of transcription control. This program is determined by the availability of the general components of the expression machinery and inevitably refers to a problem of resource reallocation, in which different parts of the genome are to use the limited expression resources.

We followed here this interpretation to present a new partition of the yeast genome through the integration of two earlier models, not connected until now (6)(11). Essential to the framework is the faculty to discriminate what it was contemplated before as a common nonspecific proportional response subjected to the same global regulation(11)-into three subclasses: invariant genes, that best follow the global program, and positive and negative genes, which were broadly defined in other studies as growth-related genes (12)(27)(28). The analysis of relative expression values is important here as it allows us to appreciate expression reallocation among partition sectors (3)(4).

We then focused on unravelling how global and specific regulation combine within this new scheme. This is largely an open question, with only a few preliminary studies in minimal bacterial systems suggesting a (somehow unexpected) minor role of TFs in the control of expression (13)(14)(15). We studied here the problem at a genomic scale using all available regulatory data to first identify invariant genes as the ones that are least regulated by TFs. The amount of regulation increases among the rest of nonspecific classes, and between these and the specific ones. Moreover, specific genes also show a stronger regulatory coherence than nonspecific genes. Coherence quantifies the similarity of expression response (to growth rate) between genes and the TFs acting on them and a strong signal suggests an operative role of the latter. In addition, among those TFs whose action is particularly coherent, we identify groups that almost exclusively regulate specific or nonspecific genes, or positive and negative genes, i.e., the functioning of the TF network is somehow segregated.

Beyond TF regulation, we discriminated two general promoter architectures. Those that are TATA enriched and shNFR/TFIID depleted (nonspecific negative and specific genes), and those that are TATA depleted and shNFR/TFIID enriched (nonspecific invariant and positive genes); features that are similarly observed in metazoans promoters (29) (Type I promoters, genes expressed in a tissue-specific manner, and Type II promoters, ubiquitously expressed genes, respectively).

That (nonspecific) positive genes are moderately controlled by TFs (like negative) but depleted in TATA box (unlike negative) could suggest certain expression features (e.g., high level of transcription, Fig. S7) and alternative modes of epigenetic regulation. Indeed, positive genes are enriched in fragile nucleosomes, which highlights the regulatory role of nucleosomal stability (16). This is supported by the particular action of GTFs on these genes [as GTFs fine-tune nucleosomal stability (19)]. In addition, we find the expression of nonspecific genes being adjusted in distinctive manner by epigenetic modifiers, with three main configurations: 1/ general activators of invariant and positive genes, 2/ remodelers working in a dual manner; repressors of negative genes and activators of positive ones, and 3/ elements acting as general repressors of negative genes.

### Global and specific regulation in a broader context

Some of the previous features match those observations related to environmental stress response genes (ESR) (30), so it is interesting to examine how this set fits into our partition. ESR genes included two complementary subsets, which are enriched in nonspecific negative and repressed genes (induced ESR genes), or nonspecific positive genes (repressed ESR genes, Methods). This confirms the suggestion of previous studies that stress response genes were not responding directly to stress but rather to the associated decrease in growth rate [but see (31)].

More generally, two models to coordinate gene expression to the available nutrients can be imagined: a *feed-forward* regulation by signaling pathways that predict growth rate in a certain environmental condition, or a *feedback* mechanism, which senses growth rate, or other related internal cell variable, and then modifies expression (32). In this context, a passive resource allocation model could explain that the global program is responding always to the environment, although indirectly (as it can only use those resources that were not consumed in the mounting of the specific response). This validates, for instance, that ribosomal genes follow the feed-forward model (33). The fine-grained structure of the nonspecific class (invariant/positive/negative) could nevertheless monitor growth rate, at least partially, with the feedback being mediated by epigenetic mechanisms (see below).

If, as suggested by Hansen and O’Shea (34), TFs can mostly transmit qualitative (presence/absence of a particular nutrient) rather than quantitative (amount of nutrient) information, how can we then explain the monotonic variation of fractional expression (with nutrient dilution in the chemostats) of the genes in the negative and positive partition components? One way is that metabolism, which is highly sensitive to the limiting nutrient (35) acts as a regulator of the epigenetic factors discussed above. Indeed, several metabolites (e.g., GlcNAc, NAD+, acetyl-CoA, alpha KG, ATP) are known to regulate transcription through interactions with enzymes involved in epigenetic modifications (36). For example, acetyl-CoA induces cell growth and proliferation by promoting the acetylation of histones at growth genes (37) (histone acetylation affects rather similarly specific and nonspecific genes, Fig. S6, what supports its potential role as a widespread mechanism). Another explanation is that the monotonic variation observed is the result of cell population shifts with growth rate, instead of changes in single-cell resource allocations. Note that these shifts cannot be attributed to the fact that slow-growing cells enlarged their G1 cell cycle phase as neither (12) nor us observed a bias in positive/negative genes with any particular phase of cell cycled genes.

In this work we have studied changes in fractional expression but not in mRNA abundances. It is known that the global program dictates that the faster a population of cells growths, the higher the promoter activity (rate of RNA synthesis) (11) or total mRNA abundance (rate of RNA synthesis and degradation) (38). We expect most (if not all) gene products to follow this (absolute) global program, with potential additional layers of regulation (which are nutrient and gene dependent) that increment or decrement mRNA levels. Expression of the genes within the invariant group best follows the absolute global program, with positive genes being slightly above and negative genes slightly below-this response (but all of them incrementing mRNA levels or promoter activities) (e.g., Fig. S1B). On the other hand, it would be interesting to quantify the degree to which single cells can present a distribution of resources that is separated from the model here discussed (39), as well as to understand the mechanisms that lead to such divergence.

In summary, although one could argue that cellular physiology can indeed determine a global transcriptional program of gene expression control, our work highlights that this program is adjusted by the integration of effective genetic and epigenetic modes of regulation. This modulation limits the prospect of “simplifying” our understanding of genome-wide expression change.

## Materials and Methods

### Promoter activity (PA) data

Keren *et al*. (11) measured the activities of ~900 S. *cerevisiae* promoters in 10 different growing conditions using a library of fluorescent reporters. For each strain in every growth condition, PA was obtained as the YFP production rate per OD per second in the window of maximal growth.

### Five-sector partition based on PA data

We computed fractional PA (f_pA_) for each growth condition as the ratio of the PA of each gene to the summed PA of all promoters (for a gene *i*, f_PA,i_ = PA_i_ / Σ PA_i_). Ratios of f_PA_s for each pair of conditions (with increasing growth rate) were also calculated. We then computed the absolute distance of these ratios to ratio 1 (i.e., same f_PA_ in both conditions), and defined as invariant genes the top 350 genes (distance closest to 0) and as activated (repressed) the bottom 50 with ratio >0 (< 0). The rest of genes with ratio >0 (<0), and both f_PA_s > 10^−4^, were designated as nonspecific positive (negative). We used the “typical” class of a gene (the most frequently occurring category that a gene presents in all pairwise growth rate changes) to functionally characterize the sectors (Table S1) what confirmed their biological significance. Minor modifications on the threshold values defining these sectors did not alter the conclusions.

### Microarray data

Brauer *et al*. (12) grew yeast cultures in chemostats under different continuous culture conditions (six different limiting nutrients each at six dilution rates) and measured mRNA abundance with two-color microarrays. Since the original reference channel for all samples corresponded to a particular glucose condition, which mixes the response of different nutrients, we reanalyzed the data without this reference by considering the red processed signal as independent channel (40), and normalizing by the corresponding sum for each case to obtain a fractional score; more specifically, the fractional expression value of gene *i* is given by f_i_ = log_10_[10^6^ (g_i_/Σg_i_], with g_i_ being the corresponding red-processed signal. SVDs were computed on this processed data.

### Five-sector partition based on microarray data

We defined as nonspecific genes those whose difference on the loadings of the 1^st^ component (a_i_’s) between two conditions is less, or equal, than three standard deviations of all gene differences (in absolute values). Genes are otherwise considered specific (activated or repressed if the difference of a_i_’s is positive or negative, respectively). Moreover, absolute values of the loadings of the 2^nd^ component (b_i_’s) were sorted to define those with smallest values (top 2500) as invariant genes, with the rest being positive or negative (determined by the sign of b_i_). To define the partition, we classify as nonspecific those genes that act as nonspecific in >8 pairwise conditions (out of 15). Nonspecific genes acting as invariant in > 3 conditions (recall that the total number is 6) are labeled as invariant. Nonspecific and not invariant genes appearing more times as positive than as negative (in all 6 conditions) are categorized as positive, and likewise for negative. Specific genes which appear more times as positive than as negative (again in all 6 conditions) are categorized as activated, and analogously for repressed. This partition is the one used for the all the regulatory analysis (Table S2 shows the connection between these sectors and those obtained with PA data). Finally, note that minor modifications on the threshold values defining these sectors did not alter the conclusions, and that the biological significance of the sectors are validated by the regulatory and functional signals observed.

### Regulatory network

We obtained regulatory data from http://yeastmine.yeastgenome.org. No microarray data is considered for the TF info; only data from different manuscripts using chromatin immunoprecipitation, chromatin immunoprecipitation-chip, chromatin immunoprecipitation-seq, combinatorial evidence, and computational combinatorial evidence for a total of 20,673 interactions with 133 TFs involved (12). We also calculated the hierarchical organization of the network (41). Bas1, Mbp1, Med6, Spt7, and Swi6 appear at the top of the hierarchy.

### Regulatory coherence

We identified the set of TFs regulating each gene and quantified the Pearson’s correlation coefficient between the expression vector (as a function of growth rate) of each TF within the set and the target gene to then take the mean. This is the (mean) regulatory coherence in a given nutrient condition. Randomizing expression vectors for each gene, 1000 times, we obtained a score of significance for each gene’s regulatory coherence. With this, we identified a list of genes displaying significant regulatory coherence. Identification of TFs acting more significantly on each partition component is computed by first measuring how often it acts on significantly coherent genes, within the five-sector partition, and then estimating a null value by randomization of the partition classes.

### Fragile nucleosome data

Nucleosome occupancy and position have been measured by analysis of MNase-digested chromatin. Recent work noted that certain nucleosomes were extremely sensitive to this digestion, and thus obtained a quantitative score of nucleosome fragility that we used for our analysis, Table S6 in (19).

### Chromatin compendium

This set includes 170 gene expression profiles for chromatin-regulation related mutations (expressed in log_2_ ratios) taken from 26 different publications (22). It covers more than 60 potential interacting chromatin modifiers such as histone acetyltransferases (HATs; the NuA4, HAT1 and SAGA complexes), histone deacetylases (HDACs; the RPD3, HDA1 and SET3 complexes), histone methyltransferases (the COMPASS complex), ATP-dependent chromatin remodelers (the SWI/SNF, SWR1, INO80, ISWI and RSC complexes), and other chromatin-affecting genes and cofactors such as Spt10, Sir proteins and the TATA-binding protein (TBP). We normalized each dataset to unit variance (21). For Fig. S6, we took absolute values to estimate the strength of the chromatin regulator effect. Note here that growth rate reduction can be connected to the impact associated to many of these deletions, so we controlled for the possible contribution of cell cycle population shifts as described (next section). This enables us to better identify expression changes due to regulation (42).

### Removal of the slow growth signature

We took the full data in (43) to obtain the slow growth profile and remove the slow growth signature in the epigenetic data following (42). In brief, the slow growth profile is obtained as the first-mode approximation of the data after SVD decomposition. To compare with the epigenetic compendium data, we chose the column of this approximation with the largest norm as the slow growth signature. The correlation with the slow growth signature is removed by transforming the epigenetic data in Gram-Schmidt fashion by subtracting from their projection onto the basis vector, given by the normalized slow growth profile.

### Statistical analysis

Null models associated to most results are obtained by randomly assigning each gene to one of the sectors, to then compute the precise statistic (e.g., mean number or regulators in Fig. 3A). We typically considered 10000 randomizations unless it is stated otherwise and show the mean and +/- 2 std deviations of the corresponding statistic (e.g., gray shading in Fig. 3A). Moreover, all P values shown in enrichment analyses (Table S1-2) are calculated using the Hypergeometric distribution with Holm-Bonferroni correction for multiple testing.

### ESR genes

There are 281 stress-induced and 585 stress-repressed genes as defined in (30)- within the set of genes delineating the five-sector partition. A subset of nonspecific negative genes and specific repressed genes corresponds to stress-induced (232 out of 2053, and 10 out of 70, respectively), while a subset of nonspecific positive genes corresponds to stress-repressed (485 out of 1914). Note that most of the features discussed in the main text associated with the five-sector partition remain when controlling for ESR genes.

## Acknowledgments

This work was supported by grant FIS2016-78781-R and the Salvador de Madariaga program (grant PRX18/00439) from the Spanish Ministerio de Economía y Competitividad (J.F.P.). We thank John C. Matese for his help with the analysis of the microarray data.

## Supplementary Information

**Figure S1.**
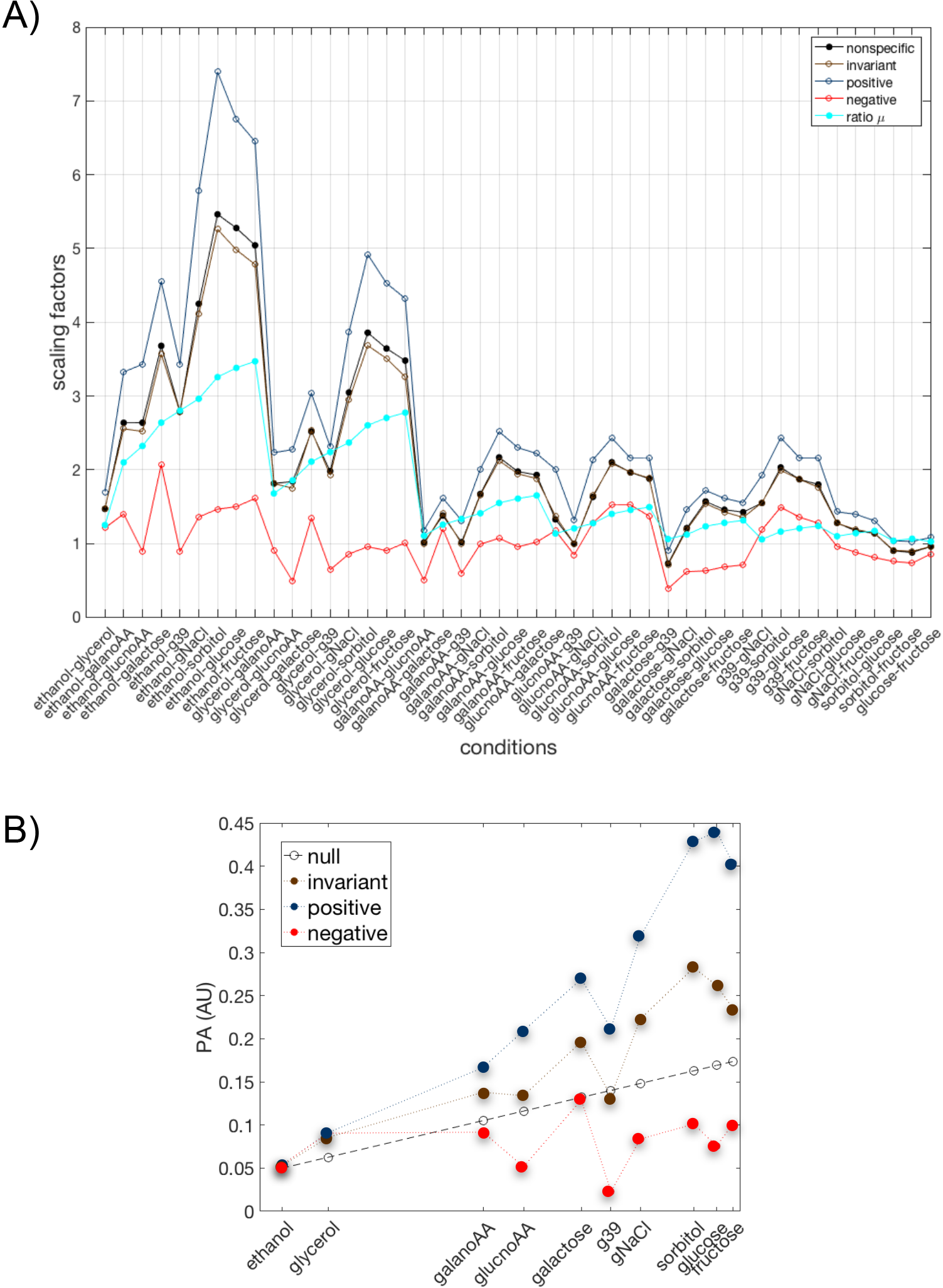
Scaling factors between pairs of conditions. **A**) Genes that follow a single proportional scaling may serve a definite cellular function according to (11). We find a single scaling that describes the change of promoter activity (PA) for different subsets of promoters according to the five-sector partition. The three classes within the nonspecific promoters (invariant, negative, positive) clearly show singular scaling. Shown also a null that corresponds to the ratio of growth rates between conditions (cyan curve). **B**) PA response of a typical invariant, positive and negative gene that corresponds to the *mrs11, rps6A* and *atp5*, respectively (conditions sorted by increasing growth rate; this is absolute PA not fractional PA). A null model of the dependence of PA with growth rate is given by the ratio of growth rates (empty circles). Gene categories within the global group clearly separate from the null.

**Figure S2.**
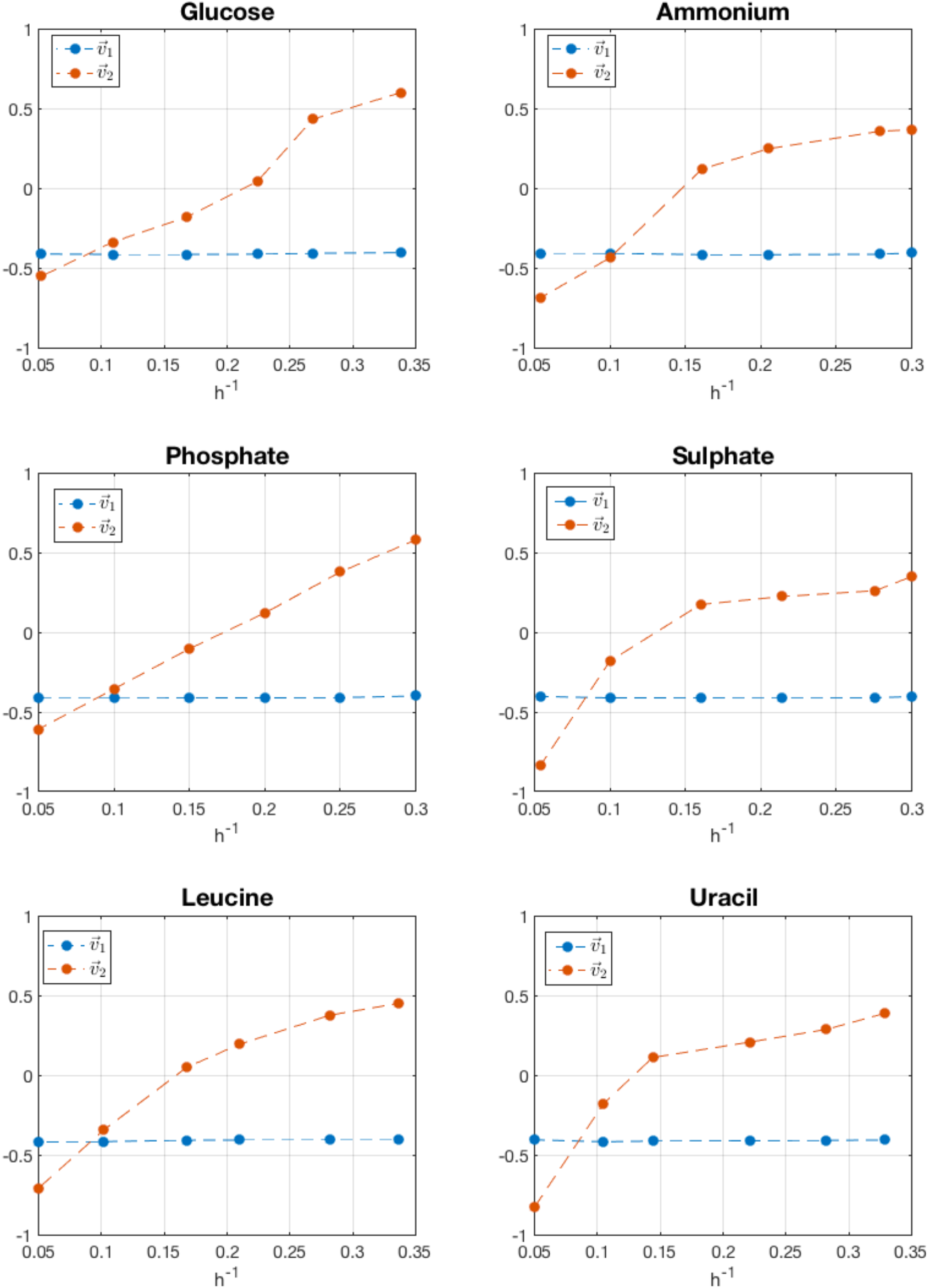
SVD components. First and second SVD components exhibited an analogous trend in all conditions what underlines a core response. As a result, expression of each gene can be approximated by the linear combination of these two components on each nutrient.

**Figure S3.**
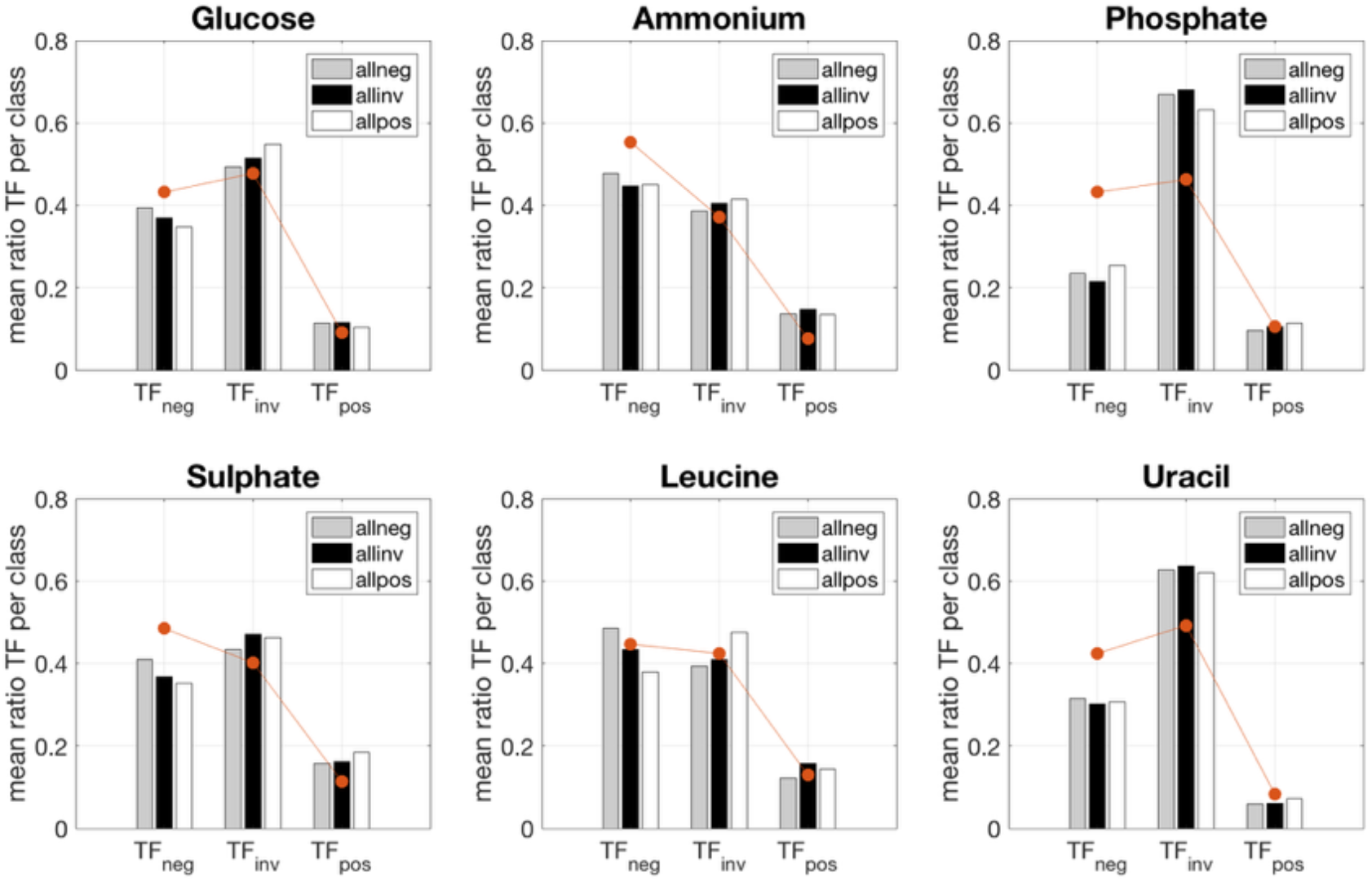
Association between class of TF and class of cognate target gene for all nutrients. Fraction of TF class (negative/invariant/positive) acting on target genes divided also with respect to growth response (negative/invariant/positive; nonspecific and specific genes were included that we denoted as allneg, etc.). Mean values of each grouping are shown in bars, while the orange curves show the distribution of each class of TF on each condition.

**Figure S4.**
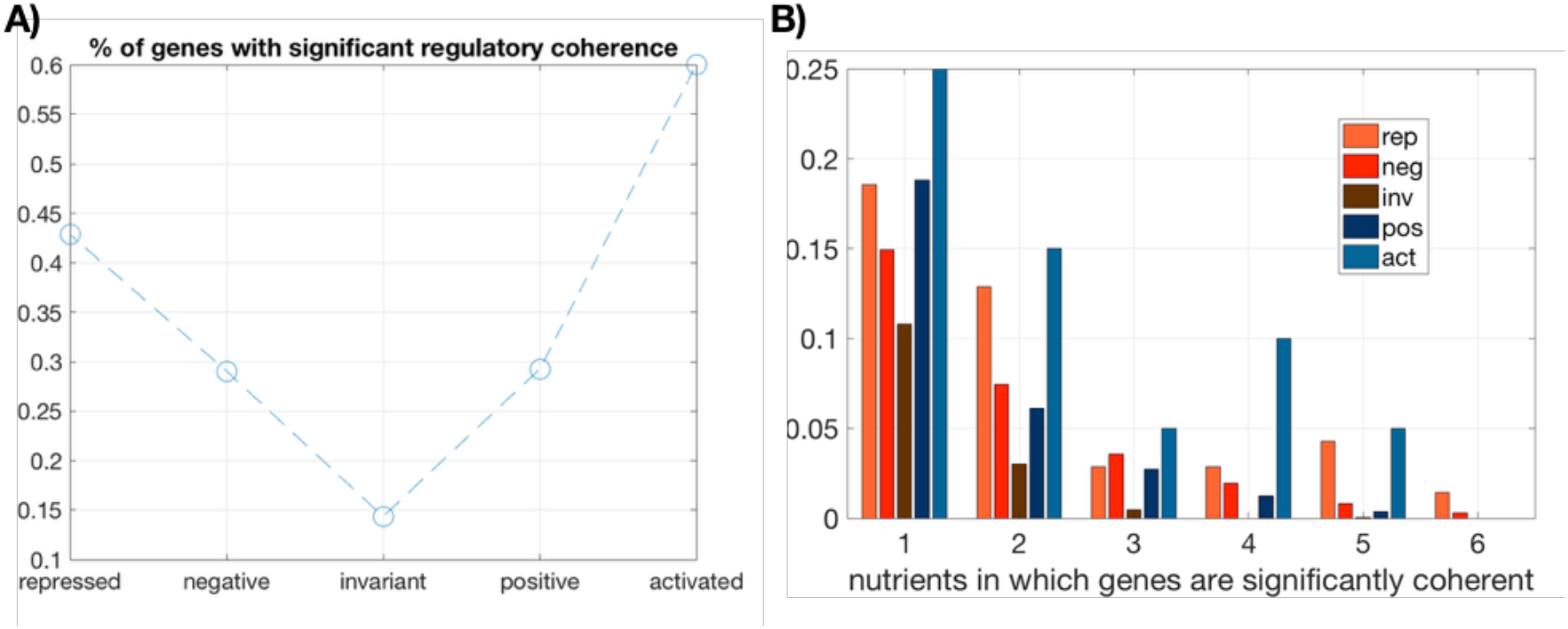
Regulatory coherence and five-sector resource allocation model. **A**) Percentage of genes within each class that exhibit a *significant* regulatory coherence with respect to a null in which classes were assigned randomly (1000 randomizations; significance implies z-scores > 2). Note that invariant genes show minimal coherence. **B**) Distribution of genes exhibiting *significant* regulatory coherence in 1 to 6 different nutrient conditions. Specific genes show more cases of genes significantly coherent in more different conditions, while invariant genes showed the opposite.

**Figure S5.**
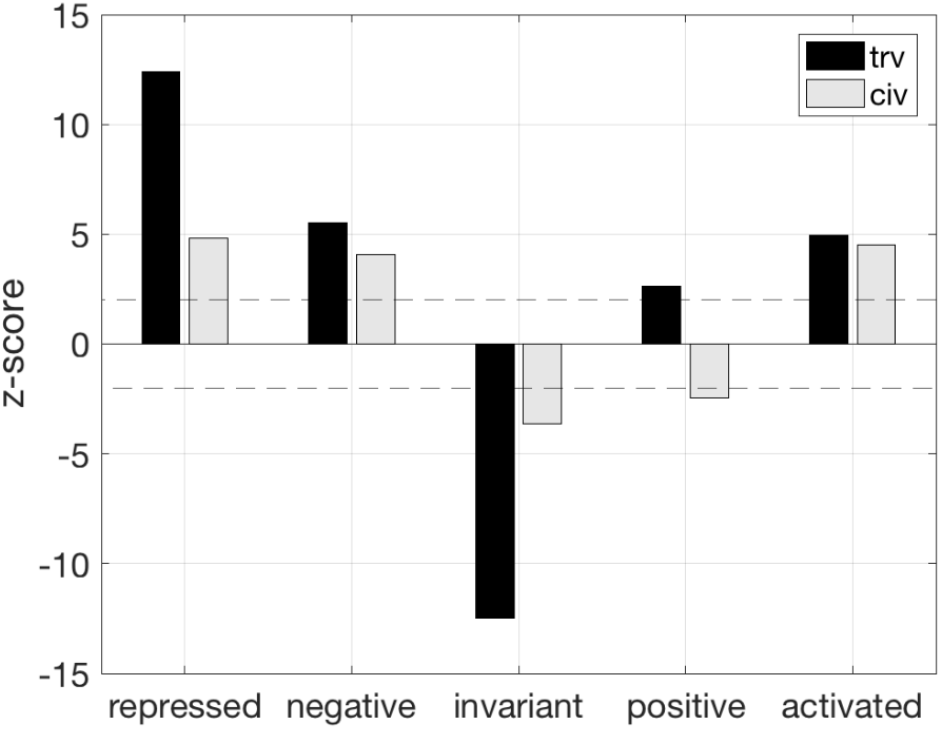
*Cis* and *trans* variability with respect to the five-sector resource allocation model. A cross between a standard laboratory yeast strain and a wild isolate allowed the computation of *cis* and *trans* effects on transcriptional variance (21). For each partition, we quantified the mean of these measures and showed the associated z-score with respect to a null by randomization; dashed line indicates z-score = +/- 2. Positive genes show dominant effects associated with *trans* variability (trv and civ denote *trans* and *cis* variability, respectively).

**Figure S6.**
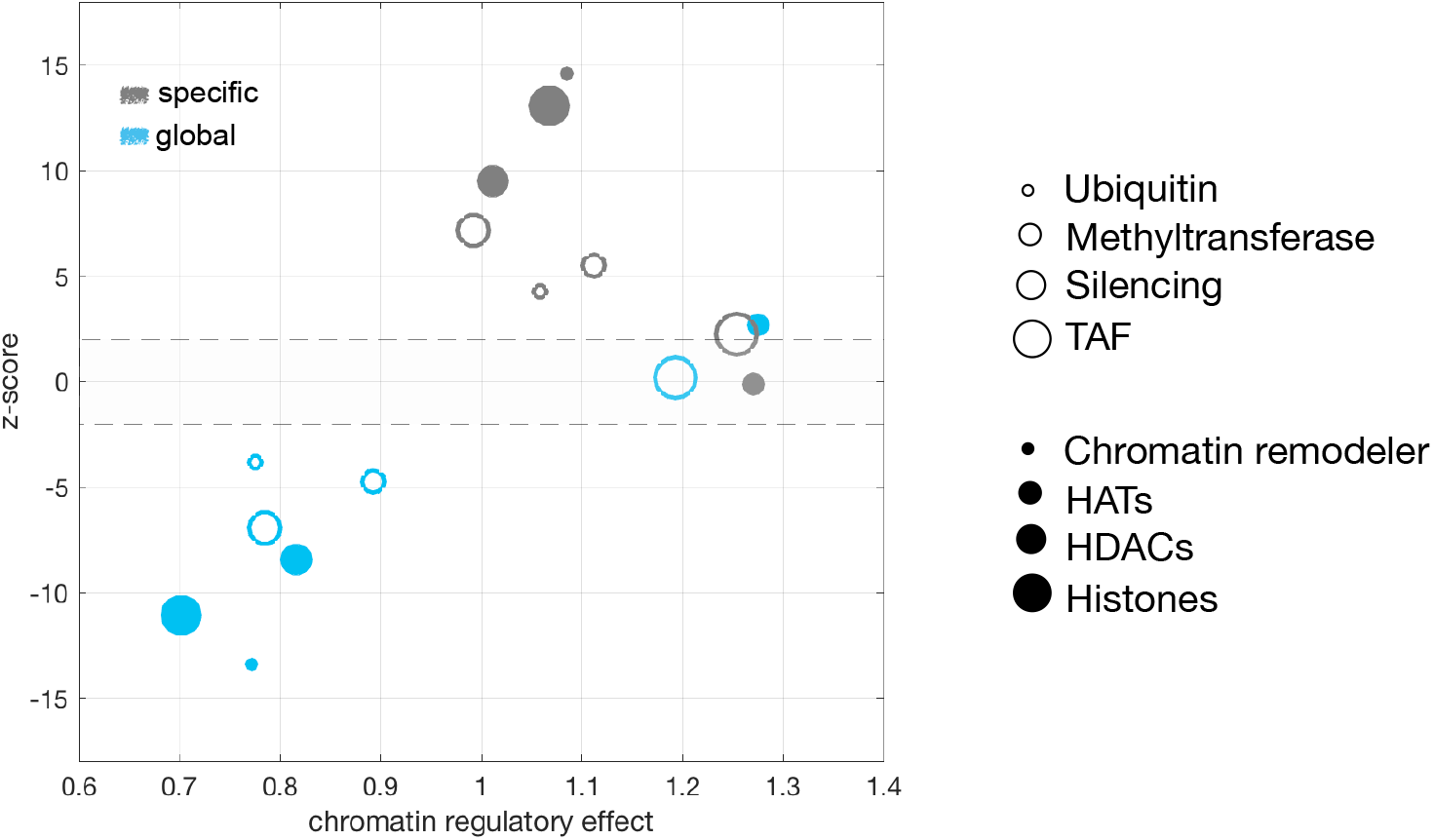
Chromatin modifiers act differentially on specific and nonspecific genes. The chromatin regulatory effect (CRE; *x* axis) quantifies change in gene expression (absolute value) due to mutations in chromatin modifiers. CRE values larger than expected by a null (z-scores > 2, obtained by randomization) are observed for most modifiers on specific genes. Each circle type corresponds to a class of epigenetic modifier (TAF: TATA-binding protein related factors; HATs: histone acetyltransferases; HDACs: histone deacetylases); *y* axis denotes z-scores and dashed lines emphasizes z-scores within +/-2 values.

**Figure S7.**
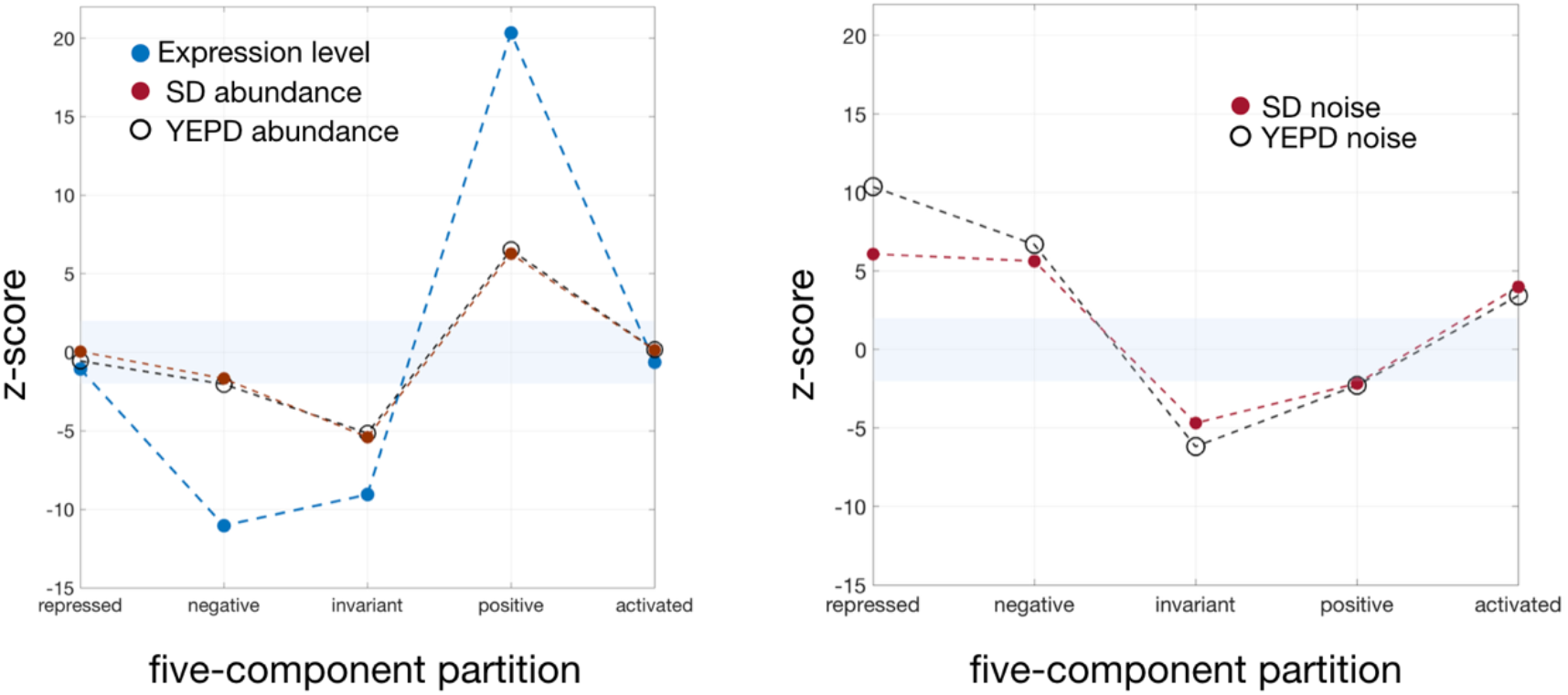
Expression abundance and noise with respect to the five-sector resource allocation model. Mean expression and protein abundance (A) and protein noise (B) with respect to the five-sector partition as compared to a null in which classes were assigned randomly (10000 randomizations; y-axis is plotting the associated z-score, shading corresponds to z-score values within a range of -/+ 2; SD/YEPD denote poor/rich growing conditions). Nonspecific and positive genes showed high expression and low noise, a signal that was associated to the presence of fragile nucleosomes in the promoter and the action of general transcription factors [both enriched in nonspecific positive genes, see main text and (24) for details on data].

